## Supplementary material for "Dual Functions of SPOP and ERG Dictate Androgen Therapy Responses in Prostate Cancer": Summplementary Fig. 1-14

#### Supplementary Figure 1

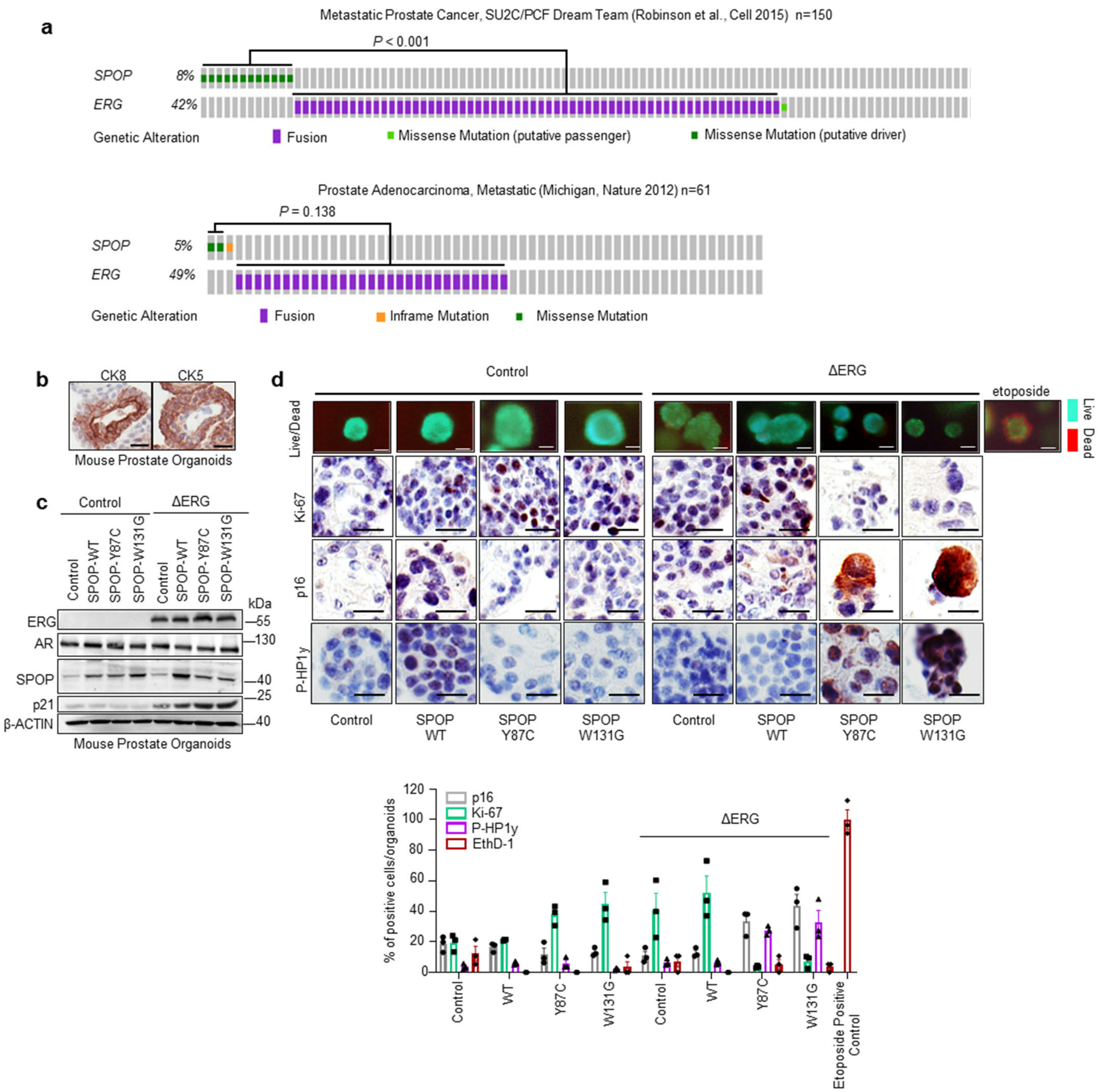


**Supplementary Figure 1. Genetic alterations in *SPOP* and *ERG* are mutually exclusive across metastatic prostate cancers and synthetic sick. a** Distribution of genetic alterations in *SPOP* and *ERG* transcription factor across 150 and 61 metastatic prostate cancer, respectively[1,2]. **b** Immunohistochemistry showing CK5 and CK8 protein expression in the mouse prostate organoids. **c** Immunoblot expression analysis of indicated proteins in mouse prostate organoids over expressing indicated SPOP mutants (MTs) and ΔERG . **d** Live and dead staining (bar 20 µm) and Ki67, p16, P-HP1y immunohistochemistry and corresponding quantification in mouse prostate organoids over expressing indicated SPOP mutants (MTs) and ΔERG**.**

**Supplementary Figure 2**

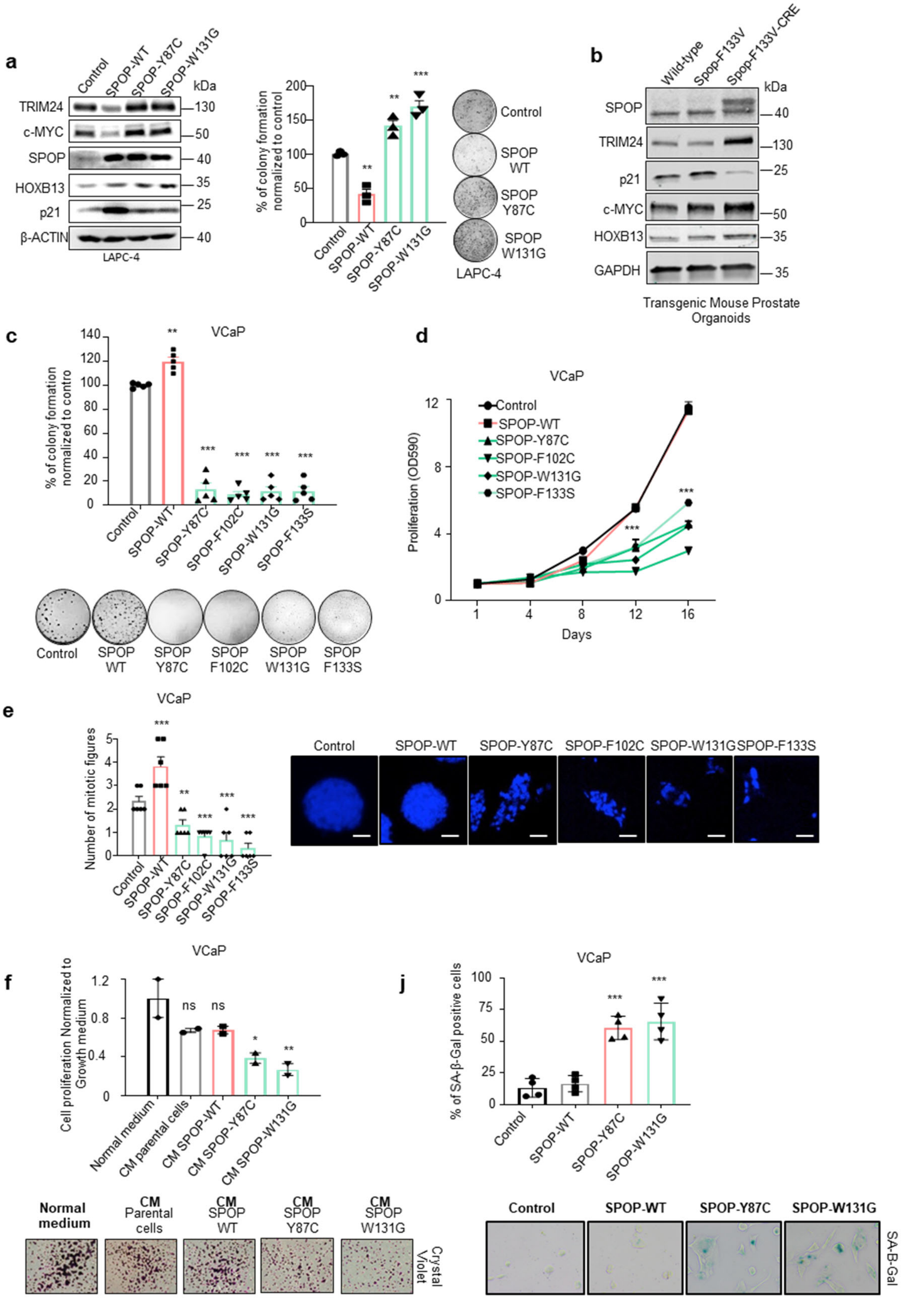

**Supplementary Figure 2. Contribution of senescence to *SPOP* and *ERG* synthetic sick relationship.** **a** 3D colony formation assay in methylcellulose of LAPC4 human prostate cancer cells over-expressing the indicated SPOP MTs and corresponding immunoblot (n=3). **b** Immunoblot of indicated proteins in Wild-type, SPOP-F133V-CRE negative and SPOP F133V-CRE positive organoids line derived from *SPOP* $^{F133V}$ transgenic mouse model. **c,d,** 3D growth in methylcellulose (n=5) and 2D proliferation assay of *TMPRSS2-ERG* positive VCaP human prostate cancer cells over-expressing the indicated SPOP MTs. **e** Corresponding mitotic count by DAPI (bar represents 100 µm). **f**, Quantification and representative images of VCaP cells proliferation assay upon normal growth medium or conditioned medium (CM) of VCaP parental cell line or VCaP cells overexpressing SPOP-WT, SPOP-Y87C or SPOP-W131G. Staining was performed after 6 days using crystal violet. CM was replaced every 3 days (n=2, independent experiment). **g** Quantification and representative images of senescence-associated β-Galactosidase (SA-B-Gal) positive cells of VCaP cells overexpressing SPOP-WT, SPOP-Y87C or SPOP-W131G (n=4). All error bars, mean + s.e.m. *P* values were determined by one-way ANOVA (**a, c, e**), two-way ANOVA (**d**) with multiple comparisons and adjusted using Benjamini-Hochberg post-test, or unpaired, two-tailed Student's t-test (**f, j**). $^{**}P < 0.01$, $^{***}P < 0.001$. Molecular weights are indicated in kilodaltons (kDa).

**Supplementary Figure 3**

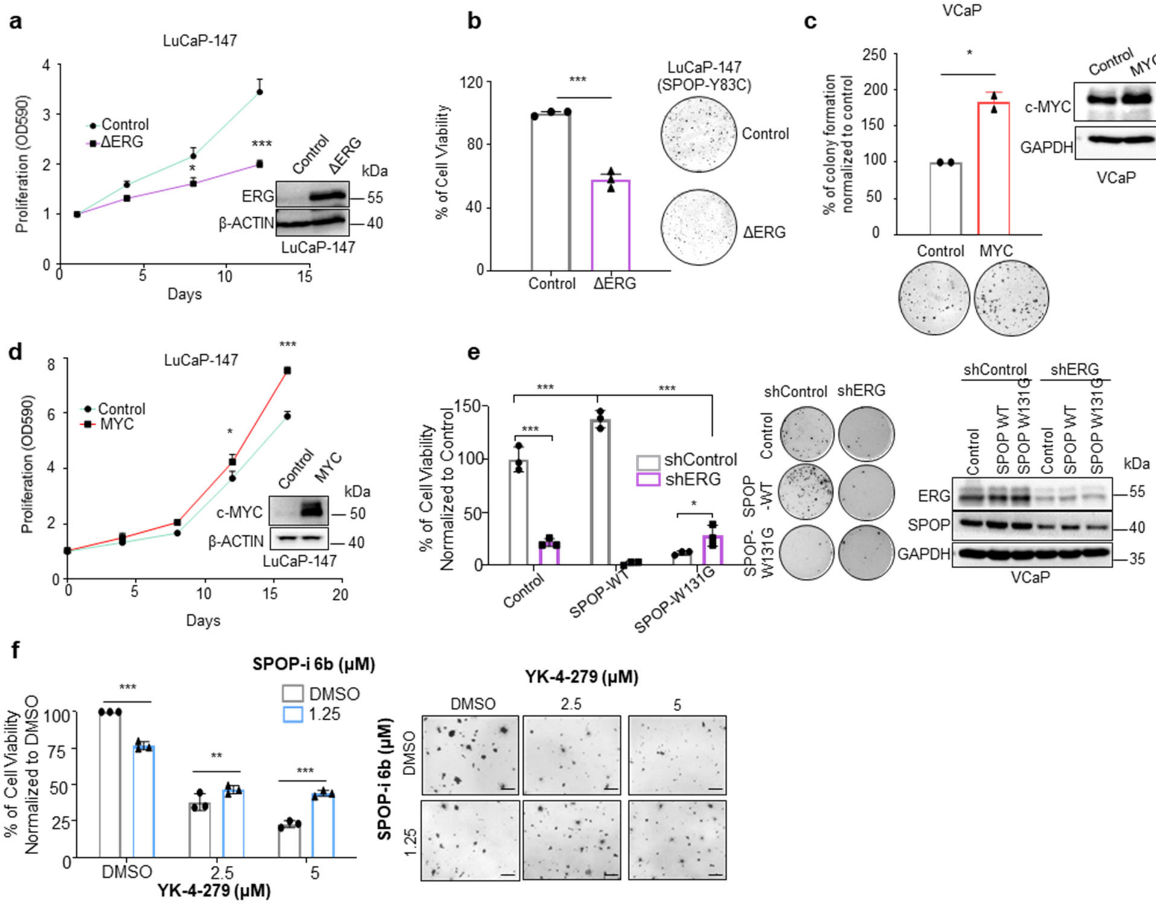


**Supplementary Figure 3. Antagonistic relationship between oncogenic activation of ERG and loss of SPOP function in prostate cancer cells. a,b** 2D (**a**) and 3D (**b**) proliferation assay of LuCAP-147 ( SPOP-Y83C) PDX cells overexpressing ΔERG and corresponding immunoblot (n=3). **c** 3D proliferation assay of VCaP cells overexpressing MYC (n=2). **d** 2D proliferation assay of LuCaP-147 overexpressing MYC (n=3) and corresponding immunoblot. **e** 3D proliferation assay of VCaP cells overexpressing SPOP-WT and SPOP-131G , with or without knockdown of ERG with a short hairpin RNA ( shERG_1) and corresponding quantification and immunoblot ( n=3). **f,** 2D proliferation assay of VCaP cells treated with SPOP-i (compound 6b) and ETS inhibitor (compound YK-4-279). Cell viability was assessed 4 days after treatment. Pictures are representative and taken after incubation with MTT reagent (bar 200 µm). All error bars, mean + s.e.m. *P* values were determined by two-way ANOVA (**a, d, e, f**) with multiple comparisons and adjusted using Benjamini-Hochberg post-test or unpaired, two-tailed Student's t-test (**b,c**) or. **P* < 0.05, ****P* < 0.001. Molecular weights are indicated in kilodaltons (kDa).

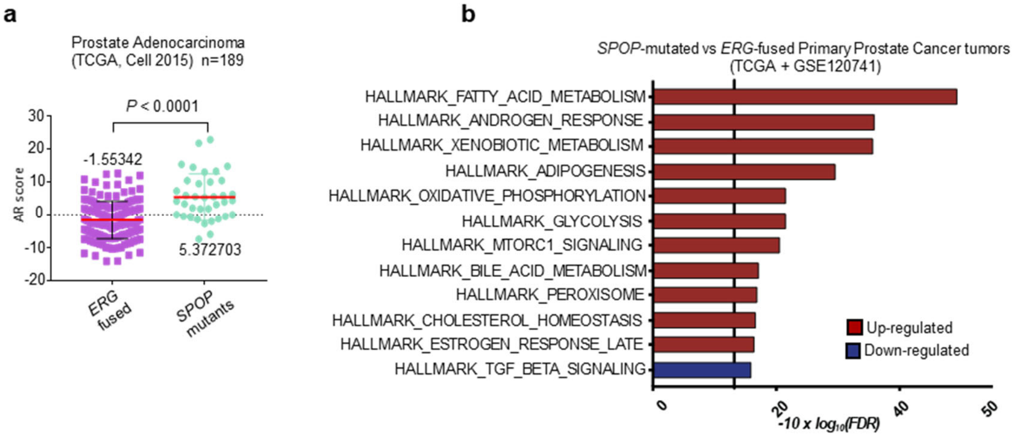


**Supplementary Figure 4. The androgen receptor signaling pathway is differentially regulated in *SPOP* mutant vs. *ERG*-fused primary prostate tumors.** **a** AR score of primary prostate tumors ERG or SPOP mutant's positive (TCGA)[3,4]. **b** Enrichment analysis of Hallmarks gene-sets performed in an integrated primary prostate cancer cohort from TCGA[2] and GSE120741[5]. FDR-adjusted p-values for individual cohorts were determined with Camera (pre-ranked) and combined user Fisher's method. Gene-sets showing opposing behavior across the two datasets were assigned a p-value of one. Association of individuals to subtypes was performed as described in Materials and Methods.

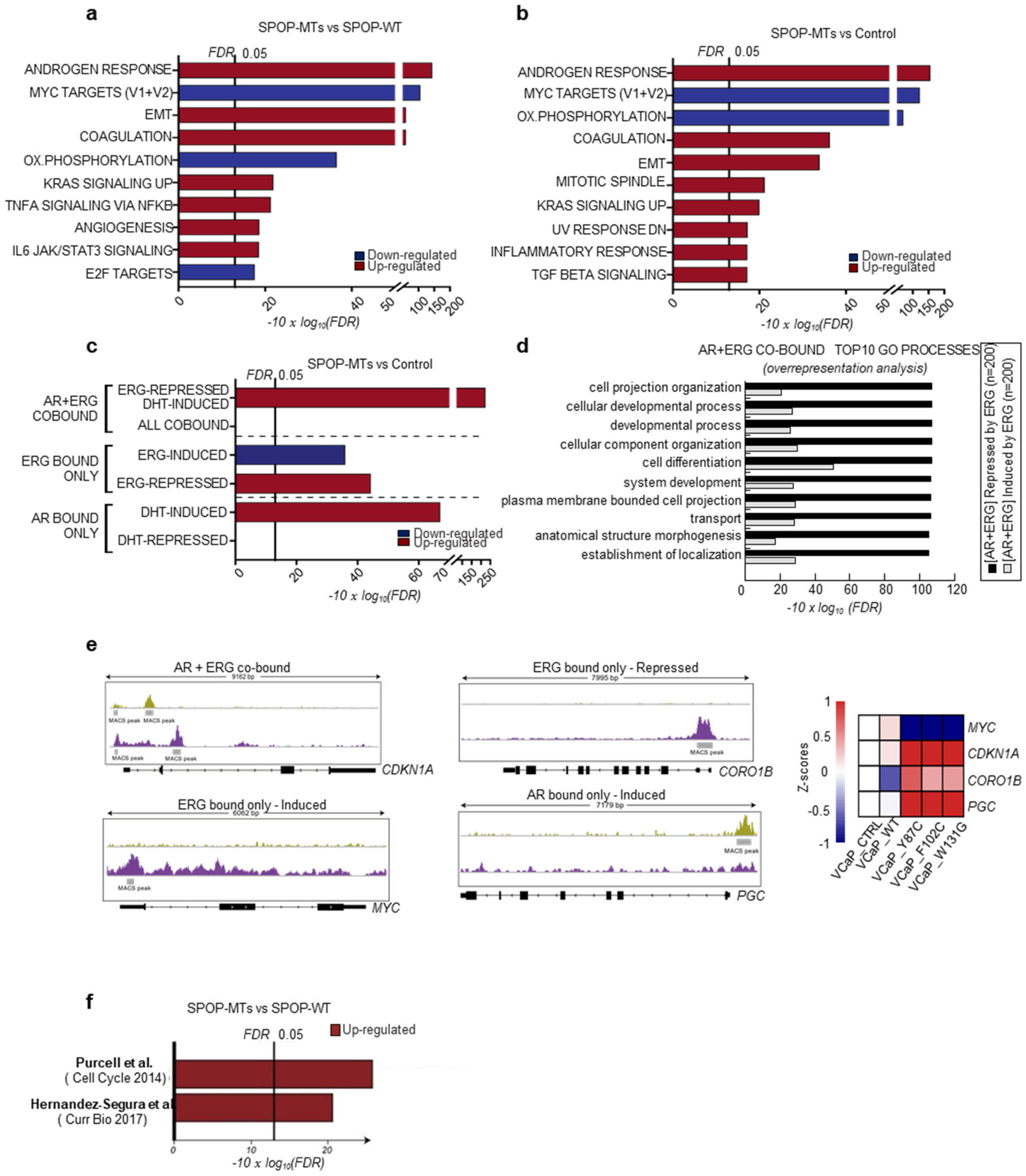


**Supplementary Figure 5. Gene expression and pathway analysis related to the synthetic sick relationship between mutant SPOP and ERG in VCaP cells. a** Gene-set enrichment analysis of SPOP-mutants (-MTs, SPOP-MTs; SPOP-Y87C, -F102C, -W131G) compared to SPOP-wild type (-WT) overexpressing VCaP cells, based on RNASeq data. Enrichments are performed on Hallmark gene-sets and FDR-adjusted p-values are determined with *Camera* (pre-ranked). **b** Gene-set enrichment analysis of SPOP-mutants (-MTs) compared to Control VCaP cells, based on RNASeq data. Enrichments are performed on Hallmark gene-sets and FDR-adjusted p-values are determined with *Camera* (pre-ranked). **c** Gene-set enrichment analysis of SPOP mutants (-MTs) overexpressing VCaP cells compared to control, based on RNASeq data. Enrichments are performed on custom gene-sets of direct androgen receptor (AR) and ERG target genes. FDR-adjusted p-values are computed with *Camera* (pre-ranked). **d** Enrichment analysis of gene ontology (GO) biological processes in the AR/ERG co-bound gene set, induced or repressed by ERG.) **e** Exemplified tracks of genes bound by AR (green) and/or ERG (violet) derived from custom ERG and AR signatures determined in VCaP cells (see Materials and Methods). Bottom: Heatmap showing mRNA expression of *MYC, CDKN1A, CORO1B* and *PGC* in control, wild-type and SPOP-mutant VCaP cells. Columns represent average expression across replicates. **f** Gene-set enrichment analysis of SPOP mutants (-MTs) overexpressing VCaP cells compared to SPOP-WT based on RNASeq data**.** Enrichments are performed on previously generated senescence gene-sets[6,7].

#### Supplementary Figure 6

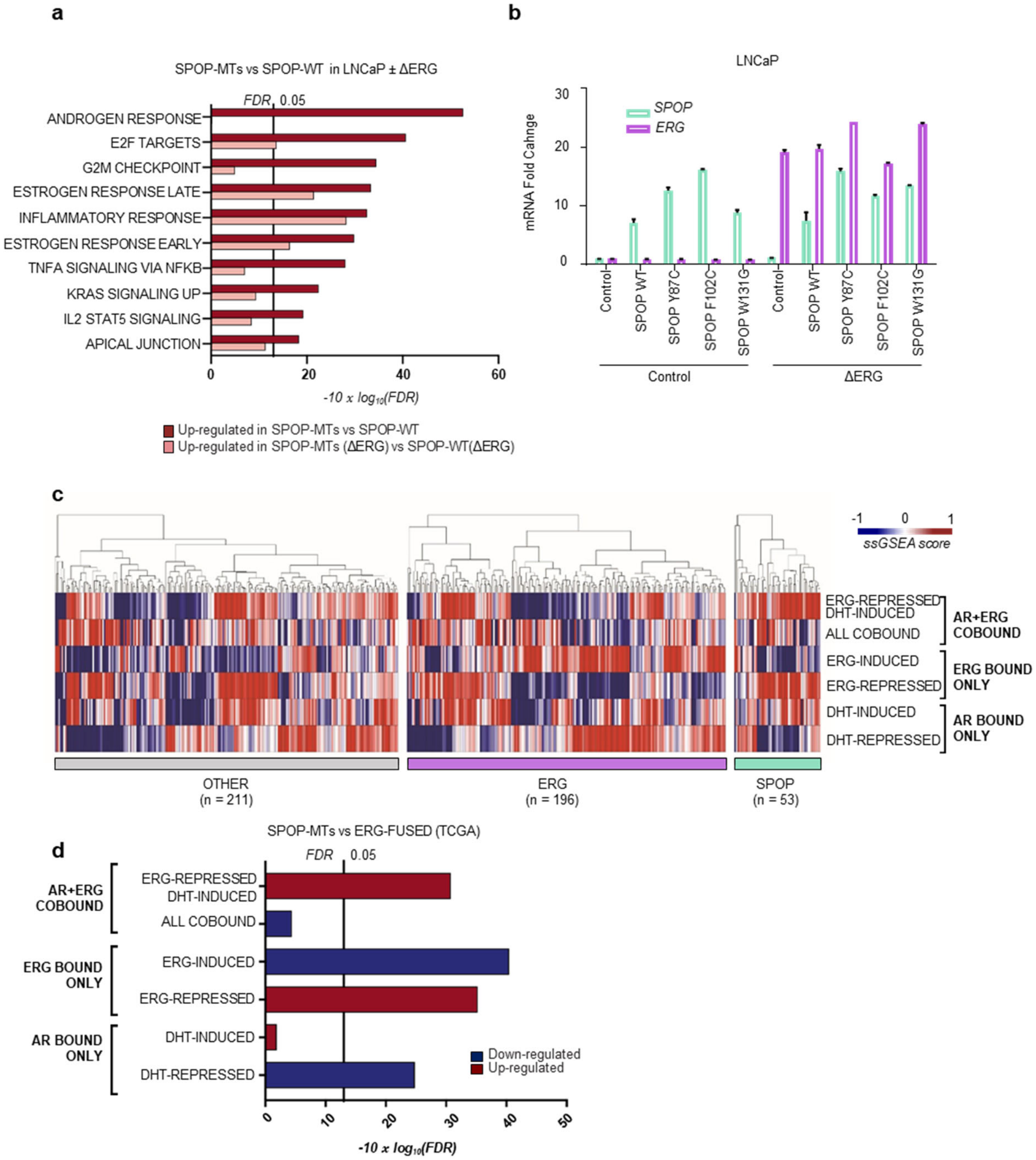


**Supplementary Figure 6. Direct AR and ERG target gene expression changes in human tumor tissues. a** Gene set enrichment analysis of LNCaP cells overexpressing either mutant (SPOP-MTs, SPOP-Y87C, or wild-type SPOP, in presence or absence of concomitant ΔERG expression. FDR-adjusted p-values were computed with Camera (pre-ranked). **b** Corresponding relative mRNA expression level of *SPOP* and *ERG* measured by qPCR. Cells were hormone-starved for 48h. **c** Heatmap of individual tumors based on single sample GSEA scores as shown in Fig. 3d. Values were scaled and centered by row. Association of individuals to subtypes was performed as described in Materials and Methods. **d** Enrichment analysis of direct AR and ERG target genes in primary prostate cancers[2]. FDR-adjusted p-values are determined with Camera (pre-ranked). Association of individuals to subtypes was performed as described in Materials and Methods.

**Supplementary Figure 7**

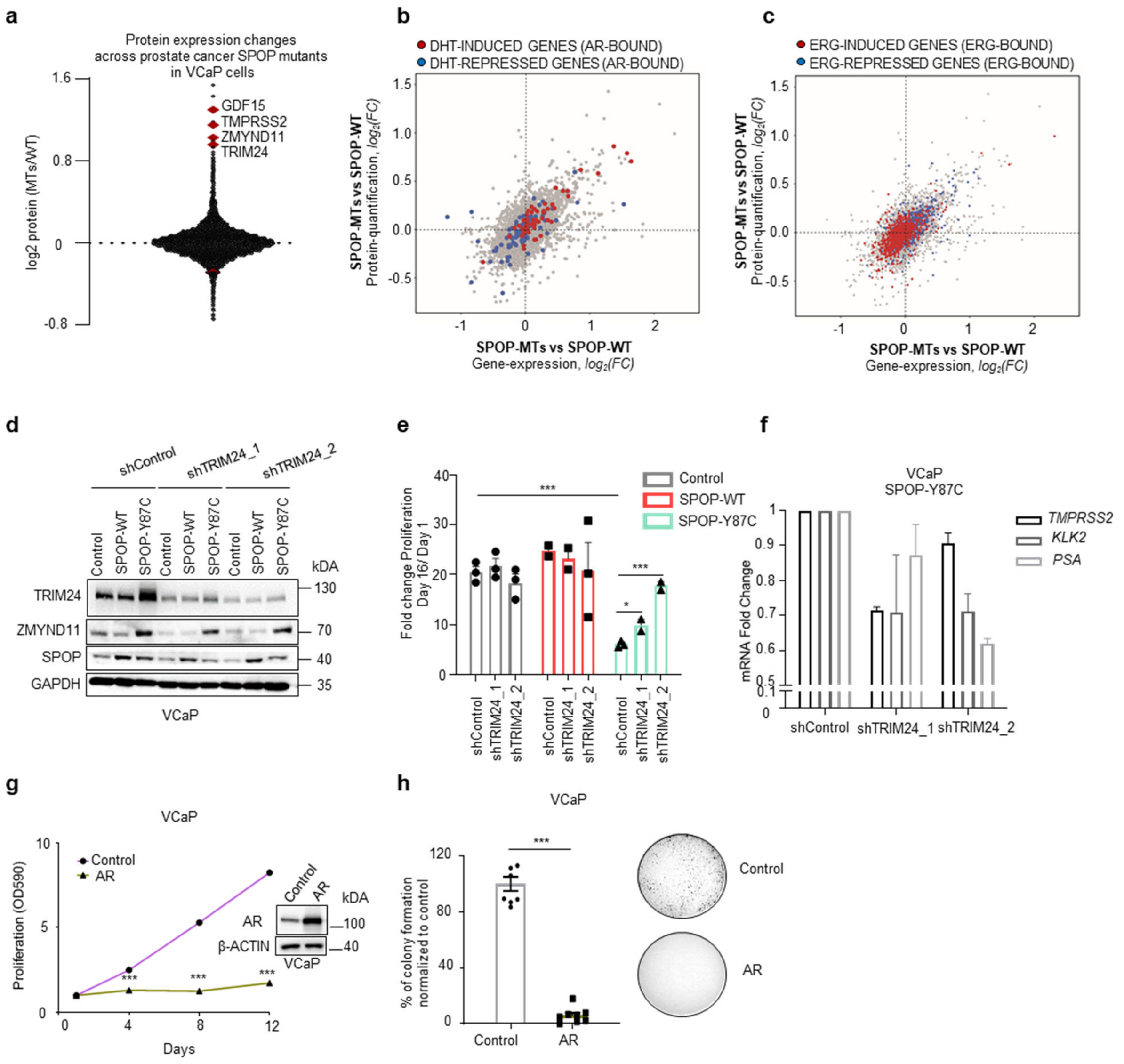


**Supplementary Figure 7. The androgen receptor (AR) signaling is implicated in the synthetic sick relationship between mutant SPOP and ERG**. **a** Protein expression changes in VCaP cells over-expressing SPOP-WT compared to SPOP mutants (average of SPOP-Y87C, -F102C, W-131G) measured by andem Mass Tag (TMT)-based quantitative mass-spectrometry (n=2, biological replicates). Top proteins being upregulated are highlighted in red (GDF15, TMPRSS2, ZMYND11, TRIM24). **b,c** Scatter plots of SPOP-MTs (average across SPOP-Y87C, -F102C, -W131G) versus SPOP-WT transcriptome and proteome expression changes in VCaP cells (n=2 biological replicates). Custom gene signatures changes are highlighted. AR bound only genes being DHT induced (red) or DHT repressed (blue) in (**b**); ERG bound only genes being ERG induced (red), or ERG repressed (blue) in (**c**). **d,e** Immunoblot of indicated proteins and 2D proliferation assay of VCaP cancer cells over-expressing the indicated SPOP mutants with and without TRIM24 knockdown using two different short hairpin RNAs (n=3). **f** AR target genes expression changes in VCaP cells overexpressing SPOP-Y87C with and without TRIM24 knockdown using two different short hairpin RNAs. **g,h** , 2D (**g**) and 3D (**h**) proliferation assay of VCaP cancer cells over-expressing

AR (n=3). All error bars, mean + s.e.m. *P* values were determined by unpaired, two-tailed Student's t-test (**h**) or two-way ANOVA (**e**,**g**) with multiple comparisons and adjusted using Benjamini-Hochberg post-test. $^*P < 0.05$, $^{***}P < 0.001$. Molecular weights are indicated in kilodaltons (kDa).

**Supplementary Figure 8**

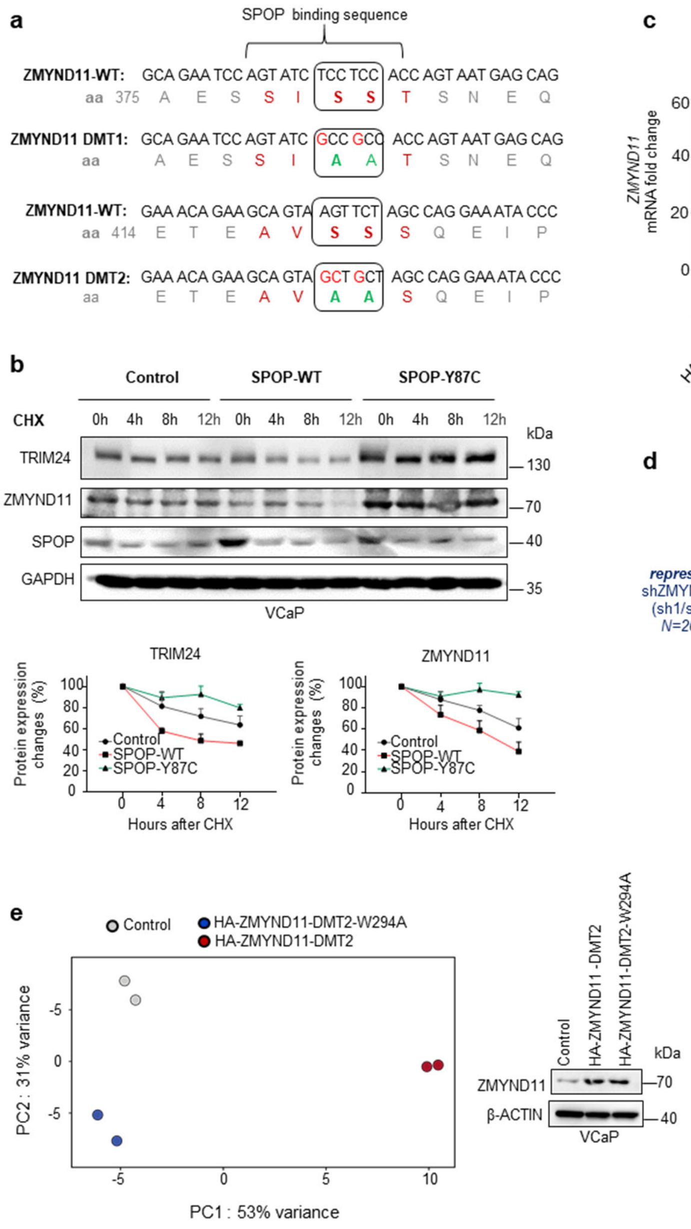


**Supplementary Figure 8. ZMYND11 is a *de novo* SPOP substrate. a** Schematic illustration of the SPOP degron sequences on ZMYND11. The degron-deficient mutants (DMT) were generated by two serine-to-alanine substitutions. **b** Immunoblots and quantification of indicated protein expression changes after treatment with cycloheximide (CHX, 100 µg/mL) in VCaP cells overexpressing the indicated SPOP species (n=2). All error bars, mean + s.e.m . Time is indicated in hours (h). **c** mRNA expression levels of VCaP cancer cells over-expressing HA-ZMNYD11-WT and derived degron-deficient mutants (DMT1/2) measured by qPCR , and corresponding 3D colony formation assay in methylcellulose (n=3). Error bars, mean ± s.e.m. *P* values were determined by one-way ANOVA with multiple comparisons and adjusted using Benjamini-Hochberg post-test. ***$P < 0.001$. **d** Chord diagram of genes transcriptionally regulated by either mutant SPOP (SPOP-MTs, SPOP-Y87C, -F102C,

-W131G) or knockdown of ZMYND11 by two different short hairpin RNAs (shZMYND11 sh1/sh2 ; shZMYND11_1 and shZMYND11_2) in VCaP cells (FDR<0.05). Strings, whose thickness is proportional to the number of shared elements, represent common genes between sets. **e** PCA-analysis based on RNA-Seq derived mRNA expression levels of the differentially expressed genes identified from the comparison between SPOP-mutant and SPOP-wild type overexpressing VCaP cells (FDR<0.05). Samples from the same experiment are shown: Controls (grey), HA-ZMYND11-DMT2 (red) HA-ZMYND11-DMT2-W294A (blue). Right: corresponding immunoblot. Molecular weights are indicated in kilodaltons (kDa).

**Supplementary Figure 9**

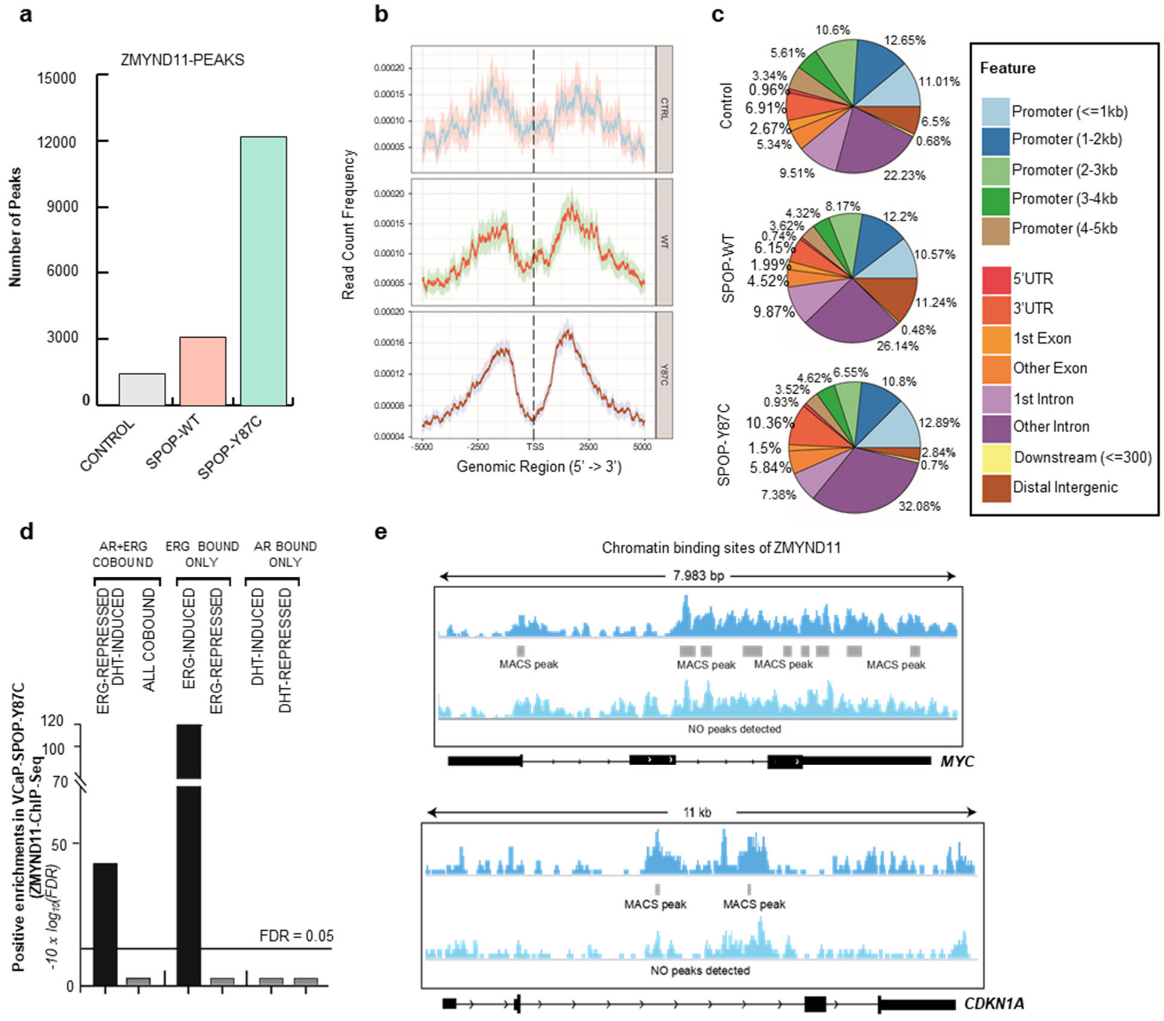


**Supplementary Figure 9. ZMYND11 influences ERG target genes transcription. a** Number of genomic ZMYND11 peaks measured by ChIP sequencing in VCaP cells overexpressing SPOP-mutant (SPOP-Y87C), SPOP-wild type (-WT) and control vector (Control). **b** Density plots representing ZMYND11 read count frequency respective to TSSs (+/- 5kb) as determined from ChIP-Sequencing experiments. Top: Control VCaP cells; Center: VCaP cells overexpressing SPOP-WT; Bottom: VCaP cells overexpressing Y87C SPOP mutation. **c** Localization of ZMYND11 binding sites, stratified according to genomic regions. **d** Enrichment analysis, performed using chipenrich[8], of genes identified by ChIP-Seq to contain ZMYND11 peaks (within 5 kb flanking each of their TSSs), performed on AR- and ERG-derived gene sets in VCaP cells overexpressing SPOP-Y87C. **e** IGV-derived screenshots representing chromatin binding sites of ZMYND11 on *MYC* (UP) and *CDKN1A* (bottom) as determined from ChIP-Seq experiments in VCaP cells overexpressing either wildtype (light pink) or mutant-SPOP (Y87C, dark green). Tracks are rescaled for their respective sequencing depth in order to be comparable.

### Supplementary Figure 10

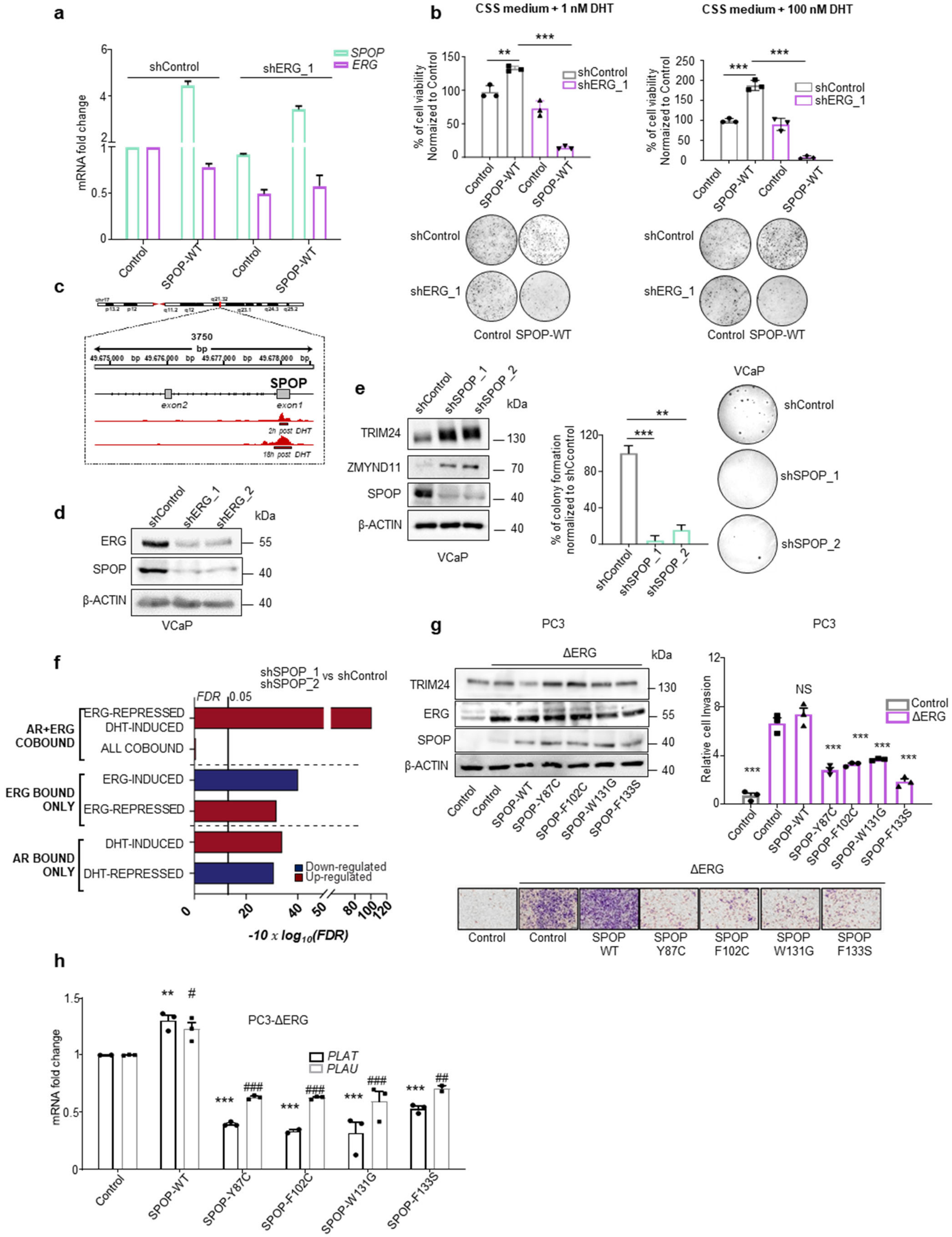

**Supplementary Figure 10. SPOP is transcriptionally up-regulated by ERG and wild type SPOP is required for ERG-mediated oncogenic phenotypes. a,b** mRNA expression of VCaP cancer cells over-expressing SPOP-WT in the context of ERG knockdown with one short hairpin RNA (**a**) and corresponding 3D proliferation assay in response to DHT treatment (**b**). Cells were plated in medium supplemented with 10% charcoal-stripped serum (CSS medium). **c**, IGV screenshot of the SPOP promoter showing ERG binding sites around exon 1[9]. **d** Knockdown of ERG with two different short hairpin RNAs in VCaP cells followed by immunoblot expression analysis of the indicated proteins. **e,** Knockdown of SPOP with two different short hairpin RNAs in VCaP cells followed by immunoblot expression analysis of the indicated proteins and corresponding 3D growth in methylcellulose. **f** Gene-set enrichment analysis of SPOP knockdown compared to shControl VCaP cells, based on RNA-Seq data. Enrichments are performed on custom gene-sets of direct androgen receptor (AR) and ERG target genes. FDR-adjusted p-values are computed with *Camera* (pre-ranked). **g** Transwell invasion assay of PC3 cells over-expressing ΔERG and indicated SPOP mutants. Corresponding protein expression changes assessed by immunoblotting (n=3). **h** Corresponding mRNA analysis of the ERG target genes *PLAU* and *PLAT* measured by qPCR. All error bars, mean + s.e.m. *P* values were determined by unpaired, two-tailed Student's t-test (**b**) , one-way ANOVA (**e, g**) or two-way ANOVA (**h**) with multiple comparisons and adjusted using Benjamini-Hochberg post-test. NS, not significant. $^{*}P < 0.05$, $^{**}P < 0.01$, $^{***}P < 0.001$; *PLAT* expression levels in Control versus each cell line ., $^{\#}P < 0.05$, $^{\#\#}P < 0.01$, $^{\#\#\#}P < 0.001$; *PLAU* expression levels in Control versus each cell line.

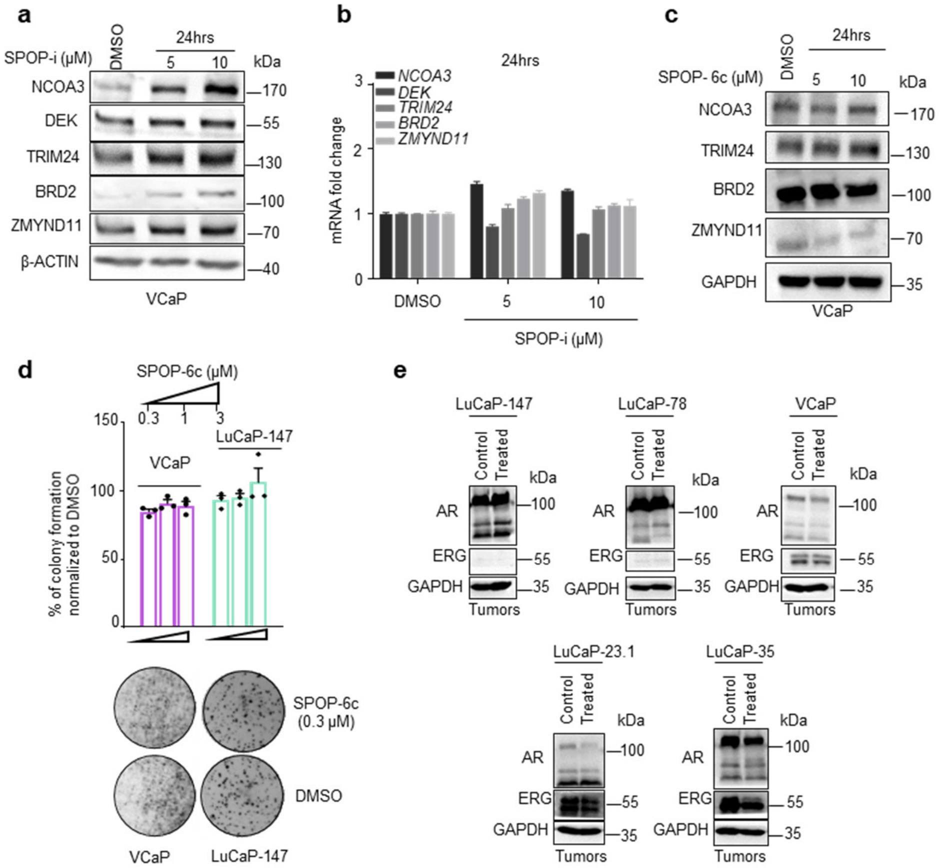


**Supplementary Figure 11. ERG-positive tumor cells are particularly sensitive to SPOP inhibition. a** Immunoblot expression analysis of indicated SPOP substrates upon treatment with SPOP inhibitor (SPOP-i, compound 6b ) in VCaP cells. **b** Corresponding mRNA expression analysis of SPOP substrates by qPCR. **c** Immunoblot expression analysis of indicated SPOP substrates upon treatment with SPOP-6c (inactive analog of compound 6b) in VCaP cells. **d** 3D colony formation assay in methylcellulose of VCaP and LuCaP-147 cells upon SPOP-6c treatment. **e** Immunoblots of indicated proteins in tumors as shown in **Figure 7b, c, d, e** and **f** respectively.

**Supplementary Figure 12**

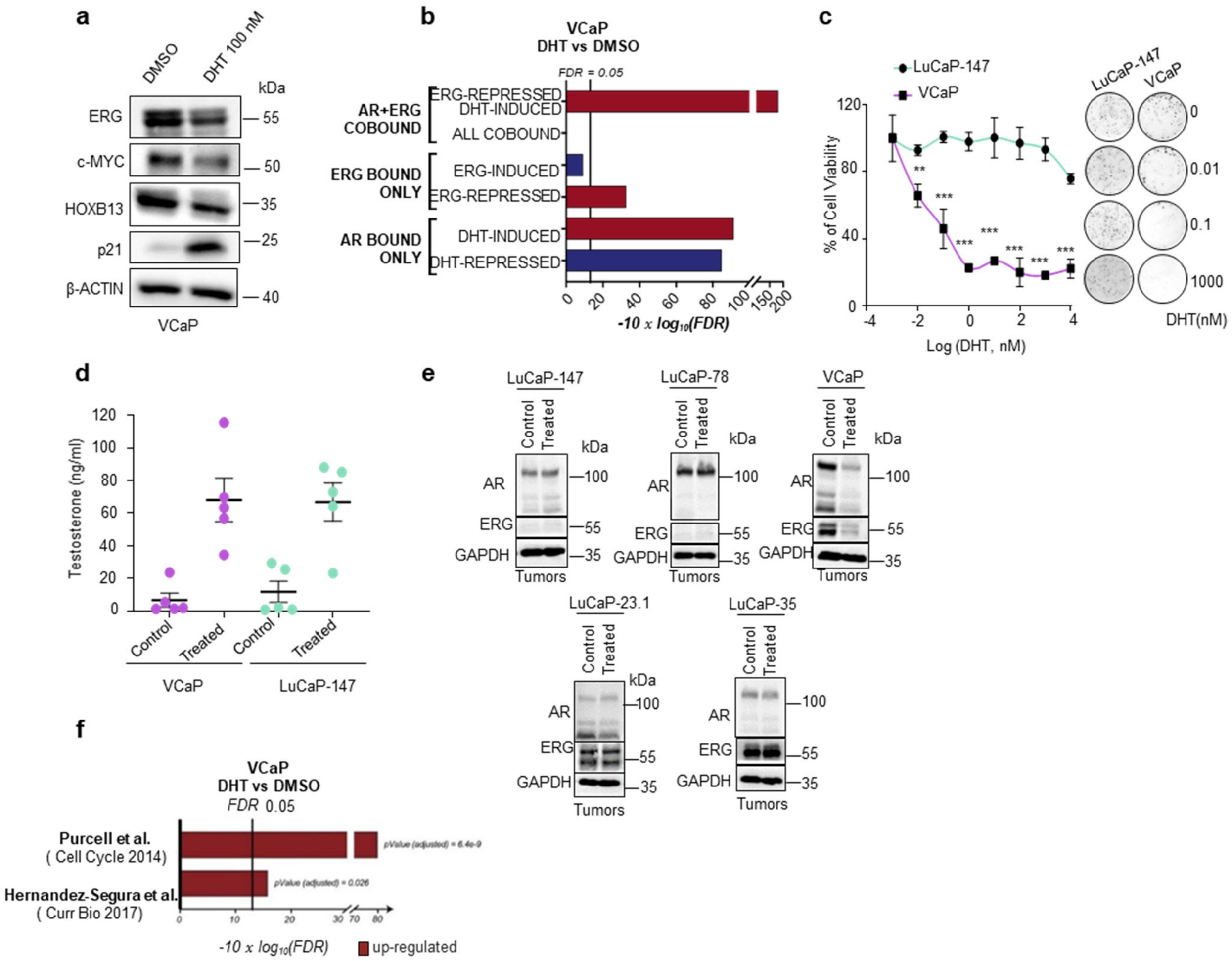


**Supplementary Figure 12. ERG and mutant SPOP trigger opposite responses to High-Dose Androgen Therapy. a** Immunoblot expression analysis of VCaP cell line treated for 7 days with DMSO or DHT (100nM). Cells were cultured in normal growth culture medium (DMEM Glutamax + 10% FBS). **b** Gene-set enrichment analysis of DHT-treated (100 nM) VCaP cells compared to control (DMSO), based on RNASeq data. Enrichments are performed on custom gene-sets of direct androgen receptor (AR) and ERG target genes. FDR-adjusted p-values are computed with Camera (pre-ranked). **c** 3D proliferation assay of VCaP (ERG positive) and LuCaP-147 (SPOP-Y83C) PDX cells in response to high doses of dihydrostestosterone (DHT). Cells were seeded in normal medium (DMEM Glutamax + 10% FBS and StemPro respectively) and treated once with corresponding DHT concentration for 10 days. **d** Testosterone levels measured in mice before and after 7 days of daily treatment**. e** Immunoblots of indicated proteins in tumors as shown in **Figure 8b, c, d, e** and **f** respectively. **f** Gene-set enrichment analysis of DHT-treated (100nM) VCaP cells compared to control (DMSO) based on RNASeq data**.** Enrichments are performed on previously generated senescence gene-sets[6,7].

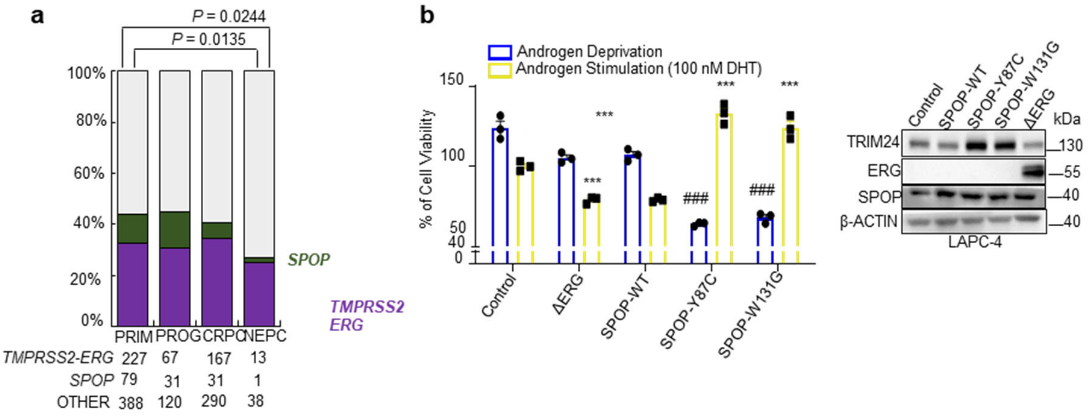


**Supplementary Figure 13. Mutant *SPOP* and *ERG*-fused tumors display different responses to androgen therapy. a** Bar plots indicating the relative frequency of *SPOP*-mutant and *TMPRSS2-ERG* positive tumors across composite primary, progressed, castration-resistant and neuroendocrine patients' cohorts (PRIM = primary; PROG = progressed; CRPC = castration resistant; NEPC = neuroendocrine)[10-12]. Statistical significance between the expected frequencies observed within primary tumors and those observed in castration resistant prostate cancer was determined by chi-squared. **b** Response to androgen deprivation or high androgen treatment (100nm DHT) of LAPC4 cells overexpressing ΔERG or indicated SPOP mutants species (Y87C, W131G) in 2D cell culture and corresponding immunoblot. Cells were cultured in CSS (charcoal-stripped serum) medium and DHT was added at the corresponding concentration. Viability was assessed after 7 days. $^{***}P < 0.001$.; Control versus each cell line under androgen stimulation. $^{\#\#\#}P < 0.001$; Control versus each cell line under androgen deprivation. All error bars, mean + s.e.m unless otherwise specified. *P* values were determined by two-way ANOVA (**b)** with multiple comparisons and adjusted using Benjamini-Hochberg post-test. NS, not significant.

#### Supplementary Figure 14

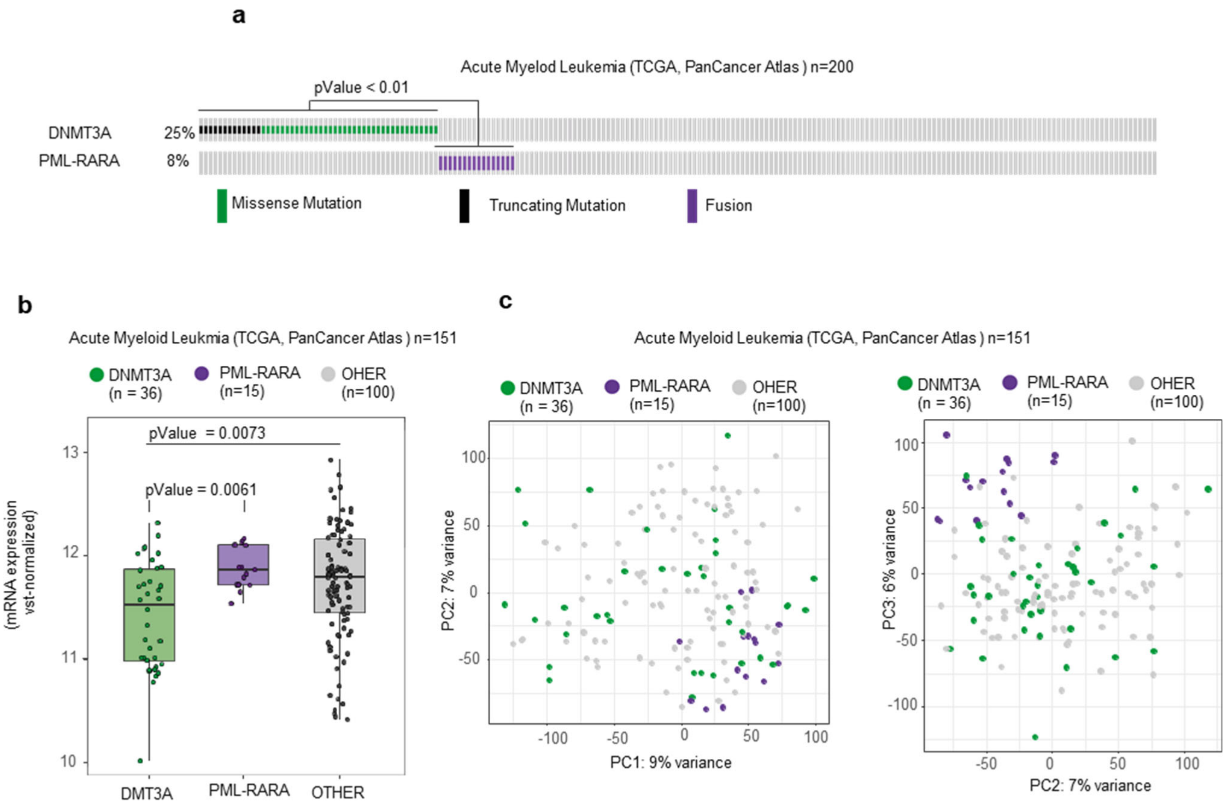


**Supplementary Figure 14.** ***SPOP* and *ERG* are part of a distinct class of antagonistic driver genes. a** Distribution of genetic alterations in *RARA* and *DNMT3* across 151 acute myeloid leukemia patients in TCGA database[13]. **b** Boxplots showing RNA-Seq based mRNA expression levels of DNMT3A among acute myeloid leukemia patients, stratified according to presence/absence of PML-RARA fusion and DNMT3A mutation status. **c** PCA-analysis based on RNA-Seq derived mRNA expression levels of the top 1000 most variable genes of the TCGA-AML cohort. RARA-fused (violet) and DNMT3A-mutant (green).
